# Fusing Mobile Phone Sensing and Brain Imaging to Assess Depression in College Students

**DOI:** 10.1101/276568

**Authors:** Jeremy F. Huckins, Alex W. daSilva, Rui Wang, Weichen Wang, Elin L. Hedlund, Eilis I. Murphy, Richard B. Lopez, Courtney Rogers, Paul E. Holtzheimer, William M. Kelley, Todd F. Heatherton, Dylan D. Wagner, James V. Haxby, Andrew T. Campbell

**Affiliations:** Department of Psychological and Brain Sciences, Dartmouth College, Hanover, NH; Department of Computer Science, Dartmouth College, Hanover, NH; National Center for PTSD, White River Junction, VT; Department of Psychiatry, Dartmouth-Hitchcock Medical Center, Lebanon, NH; Department of Psychology, Ohio State University, Columbus, OH, USA

**Keywords:** depression, mental health, smartphone, screen time, fMRI, resting-state

## Abstract

As smartphone usage has become increasingly prevalent in our society, so have rates of depression, particularly among young adults. Individual differences in smartphone usage patterns have been shown to reflect individual differences in underlying affective processes such as depression (Wang et al., 2018). In the current study, we identified a positive relationship between smartphone screen time (e.g. phone unlock duration) and resting-state functional connectivity (RSFC) between the subgenual cingulate cortex (sgCC), a brain region implicated in depression and antidepressant treatment response, and regions of the ventromedial/orbitofrontal cortex, such that increased phone usage was related to stronger connectivity between these regions. We then used this cluster to constrain subsequent analyses looking at depressive symptoms in the same cohort and observed partial replication in a separate cohort. We believe the data and analyses presented here provide relatively simplistic initial analyses which replicate and provide a first step in combining functional brain activity and smartphone usage patterns to better understand issues related to mental health. Smartphones are a prevalent part of modern life and the usage of mobile sensing data from smartphones promises to be an important tool for mental health diagnostics and neuroscience research.

## Introduction

Smartphone usage has become nearly ubiquitous in daily life at a time when depression rates are concurrently rising, particularly among college students. Smartphones contain a variety of sensors that can allow researchers to passively measure various behaviors of the phone’s user. Previous research has linked smartphone usage to self-reported depressive symptoms (Matar Boumosleh & Jaalouk, 2017; Twenge, Joiner, Rogers, & Martin, 2018; Wang et al., 2018). In parallel, depressive symptoms have been linked to brain connectivity using resting-state functional connectivity (RSFC) MRI (Greicius et al., 2007). The current manuscript has multiple goals. First, is to provide a proof-of-concept for linking passive mobile smartphone sensing technologies to brain connectivity measures that have also been linked to self-reported depressive symptoms. Second is to replicate these initial findings in a separate cohort. Third, is to identify preliminary links between a key behavior inferred from sensing (e.g. smartphone screen time) and brain connectivity metrics. Fourth, is to briefly describe a variety of methods which could be used to combine results across these various data types in the future.

### Depression assessment

Depressive disorders affect over 300 million people worldwide and have been ranked among the largest contributors to global disability since the early 1990s and currently rate as the single largest contributor to global disability (Ustün, Ayuso-Mateos, Chatterji, Mathers, & Murray, 2004; WHO depression fact sheet, 2018). Despite this, the diagnosis of depression has remained largely unchanged; further, a reliable means of identifying persons at risk of becoming depressed remains absent. Psychology, psychiatry and neuroscience have long relied upon self-reported surveys and in-person interviews to measure symptoms, diagnose mental health disorders and identify appropriate treatment strategies (Horwitz, Wakefield, & Lorenzo-Luaces, 2016). As a result of staggering costs inflicted at both individual and societal levels, clinicians and researchers set out to redefine the way mental disorders are conceptualized in hopes of creating innovative identification and prevention strategies. The aforementioned aims have been synthesized in a research framework known as RDoC (Research Domain Criteria). RDoC’s objective is to incorporate information across all planes of analysis ranging from cellular level data to person level self-report survey data to provide of a holistic picture of mental disorders (NIMH). A core principle within the RDoC framework is the notion that neuroscience will inform future psychiatric classification schemes; in other words, aid in moving towards the establishment of a neural biomarker for depression. Thus, of great importance is understanding the complete range of human behavior (and neurological functioning) from typical to atypical (Insel et al., 2010). The Patient Health Questionnaire (PHQ, with four, eight and nine question versions) is a reliable, short survey which has been validated in clinical settings and can be used to assess self-reported symptoms of depression that cause significant impairment and subjective distress (Cameron, Crawford, Lawton, & Reid, 2008; Kroenke et al., 2001; Kroenke et al., 2009), an approach in keeping within the RDoC research framework, seeking to explain individual variance in symptoms across domains, constructs, and units of analysis. Future methods to accurately diagnose depression may hold promise with the inclusion of techniques that capitalize on the passive collection of behavioral data through mobile sensors (e.g. smartphones).

### Passive sensing

Passive sensing using mobile smartphone technology allows for the assessment of daily activities by the smartphone user without continual effort on their part. This increases the frequency with which data can be collected and is less vulnerable to self-report bias, which is often a problem in prompted surveys (Ben-Zeev, Scherer, Wang, Xie, & Campbell, 2015; Rosenman, Tennekoon, & Hill, 2011). Smartphone ownership has increased steadily over the last decade, with over 75% of the US population owning one (Smith, 2017). In parallel, depression rates have increased over the last decade (Twenge et al., 2018). Prevalence of both smartphone ownership and depression rates are often reported as being higher in college-age students (Eisenberg, Hunt, & Speer, 2013; Nielsen.com, 2016). Screen time, e.g. the amount of time that the screen is unlocked and being used is a relatively simple metric to calculate. Screen time and unlock duration will be used interchangeably henceforth.

### Resting-State Functional Connectivity

Blood-oxygenation-level dependent (BOLD) functional magnetic resonance imaging (fMRI) is a non-invasive way to study activity in the human brain. Changes in BOLD signal are highly correlated with changes in neuronal activity in the local area, particularly local field potentials (Logothetis, Pauls, Augath, Trinath, & Oeltermann, 2001). Resting-state functional connectivity (RSFC) measures the relationship between the time-courses of different regions, often by using the correlation of the time-series. While connectivity across the whole brain, or “functional connectome” is fairly similar across individuals, there are small individual differences in connectivity between individuals which can be reliably observed across time. There are a variety of factors which may potentially influence RSFC, including genetics, experiences across the lifetime and current physiological and emotional state (Birn et al., 2013; Patriat et al., 2013; Poldrack et al., 2015; Richiardi et al., 2015; Shehzad et al., 2009; Sinclair et al., 2015; Zuo et al., 2014).

### Depression and neuroimaging

RSFC has been used successfully to distinguish between healthy controls and depressed individuals, even going so far as to distinguish between subtypes of depressed individuals (Berman et al., 2013; Drysdale et al., 2016; Greicius et al., 2007; Kaiser, Andrews-Hanna, Wager, & Pizzagalli, 2015). Task-based studies of self-referential processing have revealed that the sgCC is preferentially involved in processing valenced self-referential information (Moran, Macrae, Heatherton, Wyland, & Kelley, 2006; Somerville, Heatherton, & Kelley, 2006). Additionally, this region has been associated with antidepressant treatment response, and an area proximal to this has been used as a site of deep-brain stimulation for treatment-resistant depression (Holtzheimer, 2012; Mayberg et al., 2005).

### Combing RSFC and mobile smartphone passive-sensing technology

There are a wide-variety of approaches that can be taken when combining high-dimensional data from multiple modalities. We wanted to answer the following question: do smartphone sensing features previously identified as being related to depression show correlations with resting-state functional connectivity from a region previously identified to have aberrant connectivity in depressed individuals? In the current study we decided to take a targeted approach, selecting screen time with mobile smartphone (e.g. unlock duration), a feature previously shown to be linked to depressive symptoms (Twenge et al., 2018; Wang et al., 2018) and a brain area, the subgenual cingulate cortex (sgCC) which has previously been identified as having aberrant RSFC in depressed individuals, and more recently has been used as a target for deep brain stimulation for treatment resistant depression (Greicius et al., 2007; Holtzheimer, 2012; Mayberg et al., 2005). Furthermore, if there are regions identified in the passive-sensing unlock duration analysis and RSFC analysis, do these regions also show similar connectivity patterns when looking at the same correlations with brief surveys of self-reported depressive symptoms (PHQ-2, 4 and 8)? We expect that they would. Alternatively, depression may be a summation of a variety of factors and may be better understood by interrogating passive-sensing mobile technology and neuroimaging than self-reported scales. Keeping within the RDoC matrix, we assess a variety of units of analysis including brain connectivity with fMRI (physiological), passive-sensing of phone usage (behavioral) and both computer-based and phone-based depression scales (self-report).

## Methods

### Study design

In the current study two separate cohorts of first-year undergraduate students were enrolled and analyzed separately for test-retest comparison. Individuals were enrolled in three study components: neuroimaging, smartphone sensing/EMA and online surveys. Three modified versions of the PHQ-9 were used: PHQ-2/4/8. PHQ-8 is the same as PHQ-9 with the suicide ideation question removed. This question was removed before administration because the survey results are not monitored in real-time. PHQ-4 is a four question survey which includes two questions from the PHQ-8 and two from the GAD-7 as to assess both depressive and anxiety related symptoms (Kurt Kroenke, Spitzer, Williams, & Löwe, 2009). They are used because of their brief form. They may miss some of the nuances that the other inventories pick up on but have been found to have high internal reliability (Cronbach’s Alpha > 0.8) and are correlated with diagnoses of clinically relevant depression (Cameron et al., 2008; Khubchandani, Brey, Kotecki, Kleinfelder, & Anderson, 2016). PHQ-2 is used as a super-brief form of the PHQ-8 that is slightly more specific to depressive symptoms by excluding the GAD-related questions (Arroll et al., 2010).

Individuals completed an online survey to assess study eligibility (safe for MRI per Dartmouth Brain Imaging Center guidelines, no contraindications that would lead to MRI signal loss, and owned an Android or iOS smartphone compatible with StudentLife). If an individual was eligible and interested in participating in the study, she or he completed a battery of online surveys, including the PHQ-8 through REDCap (Harris et al., 2009). Individuals were then scanned during the academic term and had the StudentLife application (Wang et al., 2014) installed on their phone at or near the time of scanning. In Cohort 1, StudentLife data was collected from the time of scanning until the end of the term. In Cohort 2, StudentLife data was collected from the time of scanning and data collection is currently ongoing but the data presented here is only from their first term in college.

### StudentLife

A smartphone application, StudentLife is used in the current study to collect a variety of data about smartphone usage and mood from participants. The application is installed on a participant’s phone (iOS or Android) and collects data from the GPS, microphone, accelerometer and lock/unlock status among others. Data from StudentLife is uploaded to a secure server whenever a participant is both using WiFi and charging their phone, which they were encouraged to do daily. Data from these sensors are processed on the server to create variables that assesses the day-to-day and week-by-week impact of workload on stress, sleep, activity, mood, sociability, mental well-being and academic performance of students (Wang et al., 2014). The workflow of the current study includes data collected through StudentLife, MRI scanning sessions and self-reported surveys (Figure 1). In Cohort 1, unlock duration (phone usage) was continually sampled, providing coverage 100% of the time. This was decreased in Cohort 2 to help conserve battery usage. In Cohort 2, phones were remotely triggered every 10 minutes, sampling 1 minute every 10 minute period (minimum 10% temporal coverage), unless conversation was detected during the 1-minute sampling period, in which case sampling was extended up to 3 minutes for a maximum of 30% temporal coverage).

**Figure 1.**
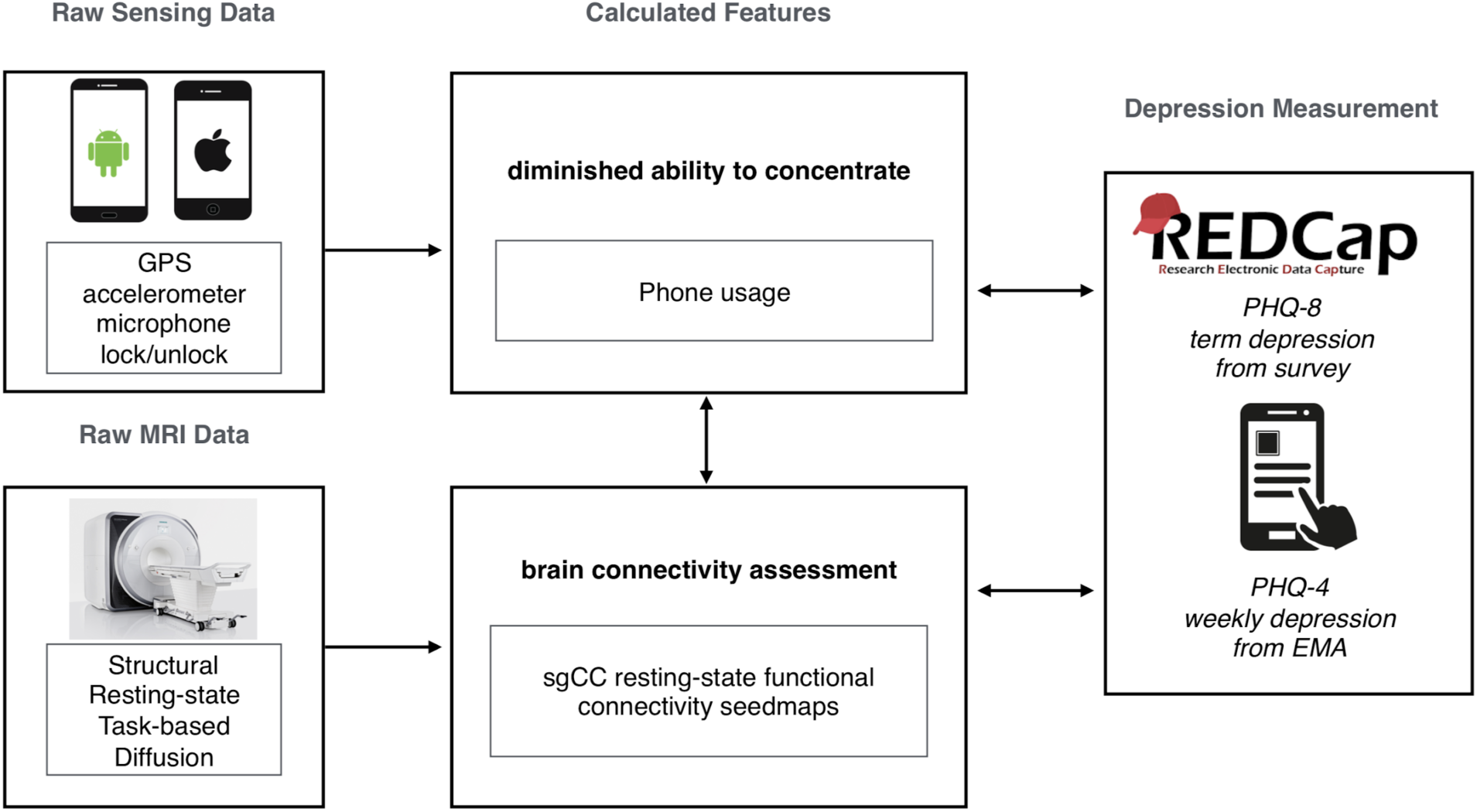
Summary graphic of the study workflow in the current study, showing raw data collection from both smartphones (StudentLife, passive sensing) and MRI (resting-state functional connectivity, sgCC seed-based analysis). Calculated features were selected based on previous research. Survey data was collected with both online (REDCap, PHQ-8) and smartphone (StudentLife, Ecological Momentary Assessments, PHQ-2/4) sources.

### Ecological Momentary Assessments

Students were prompted once a week within the StudentLife application during the term to complete a few short surveys as Ecological Momentary Assessments (EMA, one of which was PHQ-4 (Shiffman, Stone, & Hufford, 2008). In the current study we collected the PHQ-4, a modified, shorter version of the PHQ-8 which in four questions provides a glimpse of depressive and anxious symptoms (two questions related to each, with the two depression questions comprising the PHQ-2).

### Subjects

Subjects were first-year undergraduate students recruited from the Dartmouth College community. Cohort 1 included 151 subjects (94 female, mean age = 19.59, std = 1.69, range = 18-28) which were all scanned during their first year at Dartmouth and followed for the subsequent academic term. Cohort 2 included 106 subjects (75 female, mean age = 18.25, std = 0.63, range = 18-22) which were all scanned during the first academic term of their first year at Dartmouth. In Cohort 2, one subject was removed from the study for having an incompatible phone and one MRI session was stopped due to not reporting a permanent top retainer.

See Table 1 for a summary of the number of individuals included in each analysis, grouped by Cohort. Subjects were only included in each analysis if they met the minimum number of time-points for smartphone-based StudentLife data and each analysis and had resting-state functional connectivity that passed quality control (see RSFC analysis methods section below for further details). Subjects had normal or corrected-to-normal visual acuity. Each subject provided informed consent in accordance with the guidelines set by the Committee for the Protection of Human Subjects at Dartmouth College and received either course credit or monetary compensation for participating in the study.

**Table 1.**
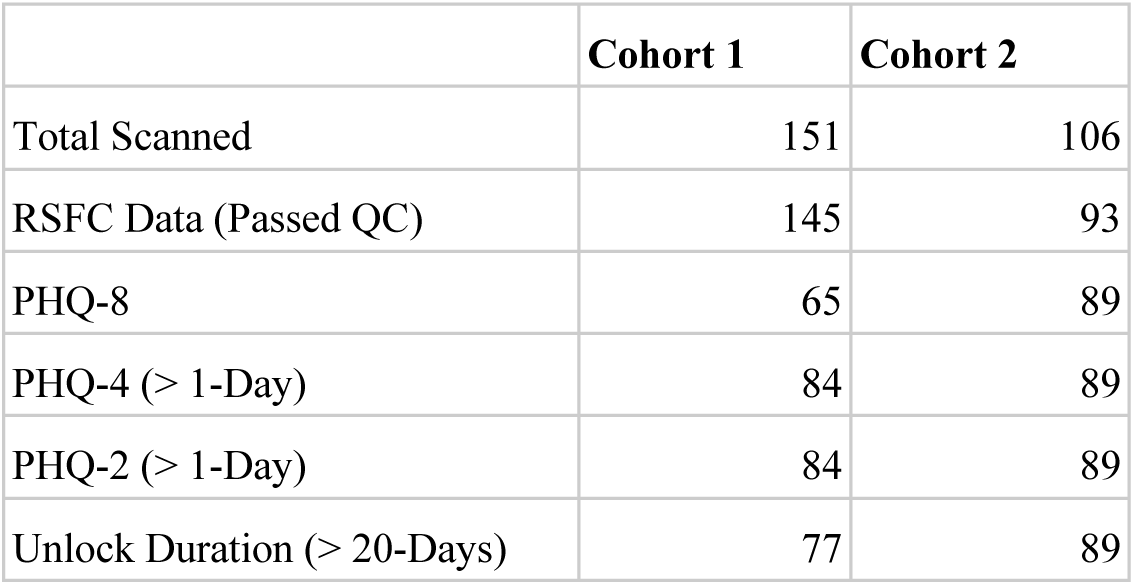
Summary of the number of subjects in each analysis.

|  | **Cohort 1** | **Cohort 2** |
| --- | --- | --- |
| Total Scanned | 151 | 106 |
| RSFC Data (Passed QC) | 145 | 93 |
| PHQ-8 | 65 | 89 |
| PHQ-4 (> 1-Day) | 84 | 89 |
| PHQ-2 (> 1-Day) | 84 | 89 |
| Unlock Duration (> 20-Days) | 77 | 89 |

### RSFC data collection

*Apparatus*

Cohort 1 imaging was performed on a Philips Intera Achieva 3-Tesla scanner (Philips Medical Systems, Bothell, WA). Cohort 2 imaging was performed on a Siemens MAGNETOM Prisma 3-Tesla scanner (Siemens Medical Solutions, Malvern, PA). Data for both cohorts was collected using a 32-channel phased array head coil. During scanning, participants viewed a white fixation cross on a black background projected on a screen positioned at the head end of the scanner bore, which participants viewed through a mirror mounted on top of the head coil.

### Cohort 1 imaging

Anatomic images were acquired using a high-resolution 3-D magnetization-prepared rapid gradient echo sequence (MP-RAGE; 160 sagittal slices; TE, 4.6 ms; TR, 9.9 ms; flip angle, 8°; voxel size, 1 x 1 x 1 mm). Resting-state functional images were collected using T2*- weighted fast field echo, echo planar functional imaging sensitive to BOLD contrast (TR= 2500 ms; TE= 35 ms; flip angle= 90°; 3 x 3 mm in-plane resolution; sense factor of 2). Functional scanning was performed in one or two runs; during each run, 240 brain volumes (36 slices, 3.5 mm slice thickness, 0.5 mm skip between slices) were acquired, allowing complete brain coverage. As such, each participant completed between 10 and 20 minutes of RSFC scanning.

### Cohort 2 imaging

Anatomic images were acquired using a high-resolution 3-D magnetization-prepared rapid gradient echo sequence (MP-RAGE; 192 sagittal slices; TE, 2.32 ms; TR, 2300 ms; flip angle, 8°; voxel size, 1 x 1 x 1 mm) with a Grappa 2 acceleration factor. Resting-state functional images were collected using T2*-weighted fast field echo, echo planar functional imaging sensitive to BOLD contrast (TR= 1190 ms; TE= 32 ms; flip angle= 63°; 2.4 x 2.4 mm in-plane resolution; SMS factor of 4). Functional scanning was performed in one or two runs; during each run, 605 volumes (46 slices, 3 mm slice thickness, no skip between slices) were acquired, allowing complete brain coverage. As such, each participant completed 12 or 24 minutes of RSFC scanning. Data for cohort 2 was processed and organized into BIDS format with datalad (Gorgolewski et al., 2016; Halchenko et al., 2017).

### RSFC analyses

All processing was performed using a standard previously published processing stream (Power et al., 2014) with two exceptions: frame-displacement (FD) threshold was set to 0.25mm (instead of 0.2mm) and 36 motion parameters (instead of 24) were used for motion regression. Functional images were preprocessed to reduce artifacts, including: (i) slice-timing correction, (ii) rigid body realignment to correct for head movement within and across runs, (iii) within-run intensity normalization such that the intensity of all voxels and volumes achieved a mode value of 1000 scale with 10 units equal to∼1% signal change, (iv) transformation to a standardized atlas space (3 mm isotropic voxels) based on (Talairach & Tournoux, 1988), (v) frame censoring, (vi) nuisance regression (excluding censored frames), (vii) interpolation, and (viii) bandpass filtering (0.009 < f < 0.08Hz) following Power et al. (2014) and using exactly the same processing stream as Huckins et al. (2018). Final correlation calculations between time-courses were calculated based upon *uncensored* frames. Preprocessing steps i-v were completed using custom scripts which call 4dfp Tools (ftp://imaging.wustl.edu/pub/raichlab/4dfp_tools/). Steps specific to resting-state functional-connectivity processing (vi-x) were completed using custom MATLAB (Version R2012b, by MathWorks, Natick, MA) scripts.

### Nuisance regressors

To control for motion, a Volterra expansion (Friston, Williams, Howard, Frackowiak, & Turner, 1996) with 36 motion parameters was used. This expansion includes motion (R), motion squared (R2), motion at the previous two frames (Rt− 1, Rt−2), and motion in the previous two frames squared (Rt −12, Rt −12). Tissue-based nuisance regressors were calculated by taking the mean signal across voxels within each of the following individual masks from FreeSurfer (http://surfer.nmr.mgh.harvard.edu) (Dale, Fischl, & Sereno, 1999; Desikan et al., 2006): an eroded (up to 4x) ventricular mask for the cerebrospinal fluid, an eroded white matter mask for the white matter signal, and a whole-brain mask for global signal. When eroded masks included no voxels, lesser erosions were progressive considered until a mask with qualifying voxels was identified. This occurred infrequently for white-matter masks while erosions of 1 were often used for CSF masks. The first derivative for each tissue regressor, as calculated by the difference from the current from to the previous frame, was also included.

### Volume censoring and data retention

Movement of the head from one volume to the next (FD) was calculated by the sum of the absolute values of the differentiated realignment values (x, y, z, pitch, roll, yaw) at each time-point (Power, Barnes, Snyder, Schlaggar, & Petersen, 2012). A frame displacement threshold of 0.25mm was used. Volumes with motion above the frame displacement threshold were identified and replaced after multiple regressions but prior to frequency filtering. Spectral decomposition of the uncensored data was performed and used to reconstitute (stage vii: interpolation) data at censored time-points. The frequency content of uncensored data was calculated with a least squares spectral analyses for non-uniformly sampled data (Mathias et al., 2004) based upon the Lomb-Scargle periodogram (Lomb, 1976). Segments of data with less than 5 contiguous volumes below the FD threshold were flagged for censoring. Functional runs were only included in the final analysis if the run contained 50 or more uncensored frames. Only subjects with at least 5 minutes of uncensored data across runs were included in the current study. Consistent with Power et al. (2014), only uncensored volumes were used when calculating temporal correlations.

### Neurosynth analysis and subgenual cingulate cortex seedmaps

To identify an unbiased subgenual cingulate cortex (sgCC) seed to create voxelwise functional seed maps, an automated meta-analysis was performed using Neurosynth for the term “subgenual” (Yarkoni, Poldrack, Nichols, Van Essen, & Wager, 2011). Subgenual cingulate cortex seed maps were created from a 4mm spherical seed placed at 0, 25, -10 (MNI coordinates), which was the peak of the term “subgenual” as of February 17th, 2017 and are centered around BA 25. The mean time-course from this seed was correlated with the time-course from every voxel within the brain. These seed maps, i.e. maps of resting-state connectivity from the subgenual region, were produced for each individual that passed quality control (more than 5 minutes of uncensored frames, see above for more details).

### Combining data

Since the version of the StudentLife application used in the current study generates 182 features automatically, and with RSFC it is possible to generate thousands of features, it is necessary to minimize the number of features compared given the relatively small size of the Cohorts (N<100). To minimize the number of features inspected, unlock duration was the only feature inspected given its simplicity to calculate and previously-identified relationship with PHQ-8 (Wang et al., 2018). While many features were identified, we specifically chose unlock duration (e.g. screen time) as a simple feature both to calculate and to conceptualize as it can be considered similar to total phone screen time.

For all surveys analyzed here, one time-point was sufficient for a subject to be included in the current analyses. If there were multiple responses to ecological momentary assessments (EMAs, e.g. surveys prompted by the application) over the course of the term those responses were averaged. Individuals were included in the passive sensing unlock duration analysis if they had 20 days of quality data with more than 16 hours of quality unlock duration data for each day included.

### Group analyses and statistics

SgCC seedmaps from Cohort 1 were correlated with unlock duration sampled from smartphone usage with the StudentLife application. For each analysis, the degrees of freedom was N-2, with N being the number of subjects which is listed in Table 1. Results from the unlock duration and sgCC correlational analysis from Cohort 1 were volume corrected to account for multiple comparisons using AFNI’s 3dClustSim ACF function. Results from the sgCC/unlock duration analysis were used to restrict the regions investigated in further analyses. Given the proof-of-concept and exploratory nature of the current work, clusters are marked as having passed volume-correction or not.

### Visualization

All results were transformed into MNI space (Montreal Neurological Institute) and mapped onto the Conte69 template for volume-based slices or inflated surfaces for visualization (Van Essen, Glasser, Dierker, Harwell, & Coalson, 2012). Group results were viewed in Connectome Workbench Version 1.1.1 (Marcus, Fotenos, Csernansky, Morris, & Buckner, 2010).

## Results

### Unlock duration correlated with sgCC connectivity

In Cohort 1 exploratory whole-brain analyses of the correlation between unlock duration and sgCC seedmaps identified a large cluster (584 voxels, 15,768mm^3^) in the ventromedial prefrontal cortex with a positive linear relationship (Figure 2, S1A). This cluster extended from the ventral striatum to medial frontal orbitofrontal cortex and dorsally to medial prefrontal cortex. Information about subpeaks within this cluster can be found in Table 2. To determine if these results replicated in Cohort 2, the cluster identified in Cohort 1 was used as a mask and voxels which showed a significant positive relationship between unlock duration and sgCC connectivity in Cohort 2 were identified. This analysis identified a cluster with the peak located at -6, 51 -18 (MNI coordinates, peak T = 2.94, voxel extent = 42, volume-corrected to p<0.05) (Figure S1B).

**Figure 2.**
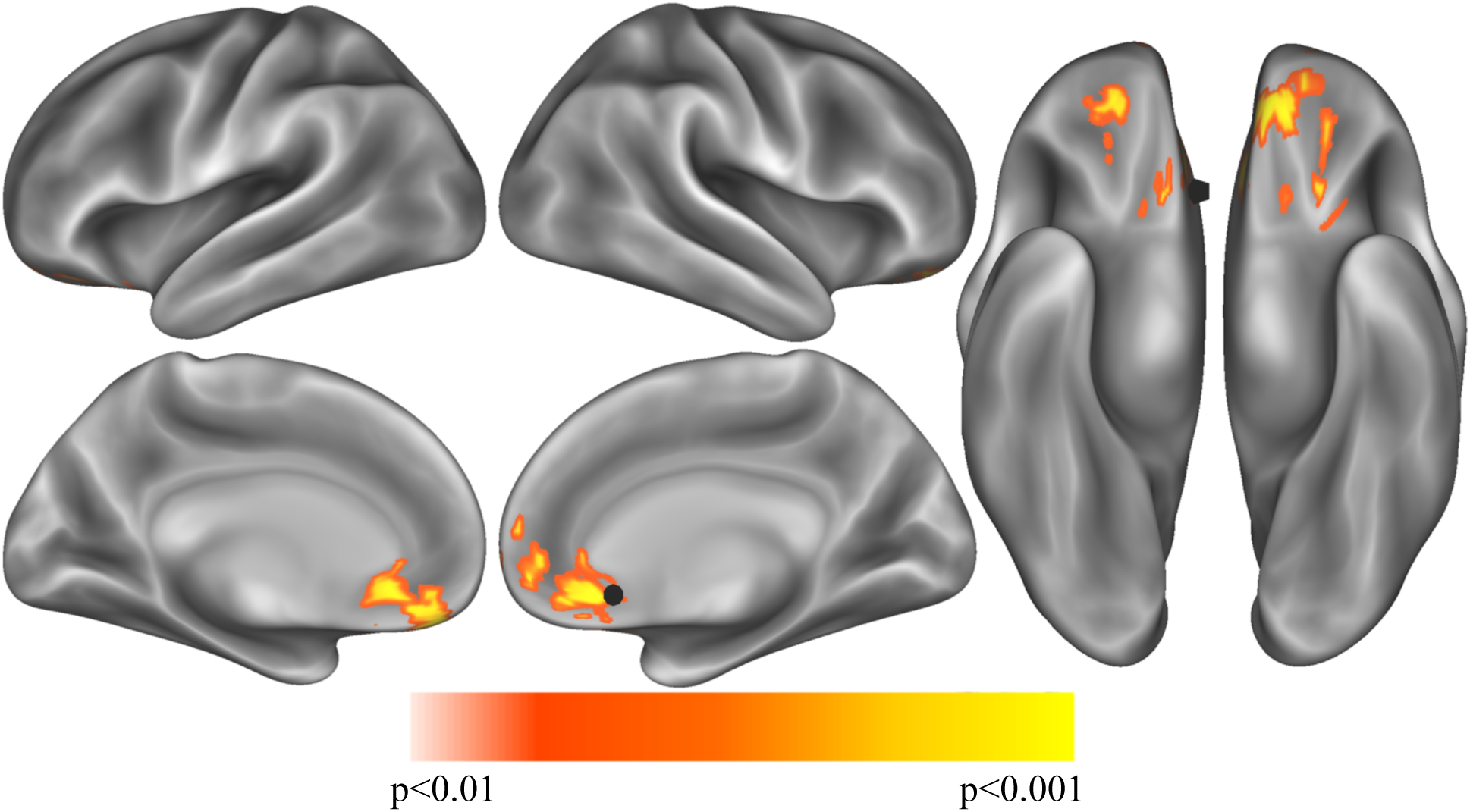
Exploratory analysis correlation sgCC RSFC seedmaps correlated with mean unlock duration identified a cluster with a positive relationship to unlock duration in the ventromedial prefrontal cortex (p<0.01, volume corrected using ACF to p<0.001) shown on inflated lateral (top left), medial (bottom left) and ventral (right) cortical surfaces. The sgCC seed is represented as a black 10mm sphere, larger than the 4mm sphere used to create the seedmaps for visualization purposes.

**Table 2.**
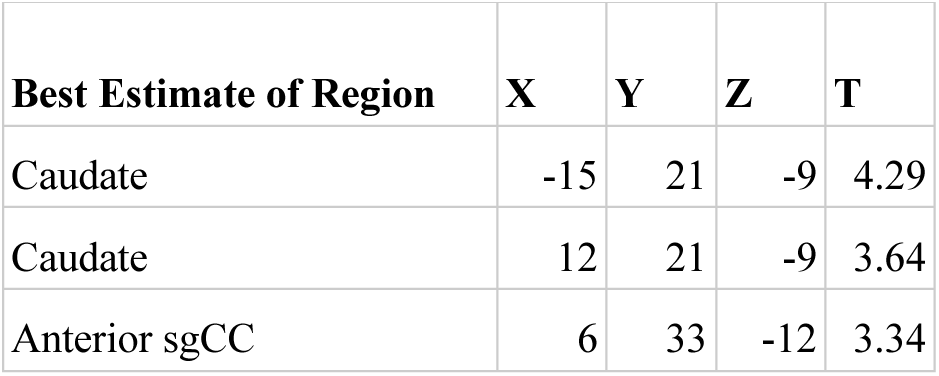
Exploratory analysis correlation sgCC RSFC seedmaps correlated with mean unlock duration (smartphone screen time) identified one cluster in the ventromedial prefrontal cortex (p<0.01, volume corrected using AFNI’s ACF to p<0.001, k>449, voxel extent=548). Peaks were identified with xjview 9.6, showing 3 maximia within this cluster, at least 8mm apart.

| Best Estimate of Region | X | Y | Z | T |
| --- | --- | --- | --- | --- |
| Caudate | -15 | 21 | -9 | 4.29 |
| Caudate | 12 | 21 | -9 | 3.64 |
| Anterior sgCC | 6 | 33 | -12 | 3.34 |

### Self-reported depression symptoms correlated with sgCC connectivity

Previous research (Wang and colleagues, 2018) identified a relationship between depressive symptoms and unlock duration. To determine if depressive symptoms and unlock duration had overlap in the brain connectivity (seed based subgenual RSFC) regressions for both computer-based pre-screening (PHQ-8), phone based post-scanning (PHQ-2/4 as EMA) were performed. Results from each of these analyses were masked with the cluster identified in Cohort 1’s sgCC/unlock duration analysis.

PHQ-8 computer-based surveys correlated with sgCC connectivity maps identified clusters with a positive relationship with sgCC connectivity in both Cohorts and identified a cluster which overlapped between the two. Cohort 1 revealed one cluster at which passed volume-correction -21, 42, -12 (peak T = 3.19, voxel extent = 63, volume corrected to p < 0.05), 24, 51, -9 (peak T = 2.55, voxel extent = 15, did not pass volume correction) (Figure 3, Table 3). In the PHQ-8 analysis of Cohort 2, results were further masked by the cluster which passed volume-correction in the Cohort 1 PHQ-8 analysis (63 voxels), identifying 1 significant cluster in Cohort 2, located at -15, 33, -12 (peak T = 2.98, voxel extent = 8, volume corrected to p < 0.05). In addition to identifying a cluster with overlap between the both Cohorts for the PHQ-8 analysis, qualitative visual inspection suggests proximal cortical regions in both cohorts meeting a voxelwise threshold of p<0.05, with regions proximal to the mask having overlap at a threshold of p < 0.05 and increased overlap, including right OFC at a more liberal threshold of p < 0.1.

**Figure 3.**
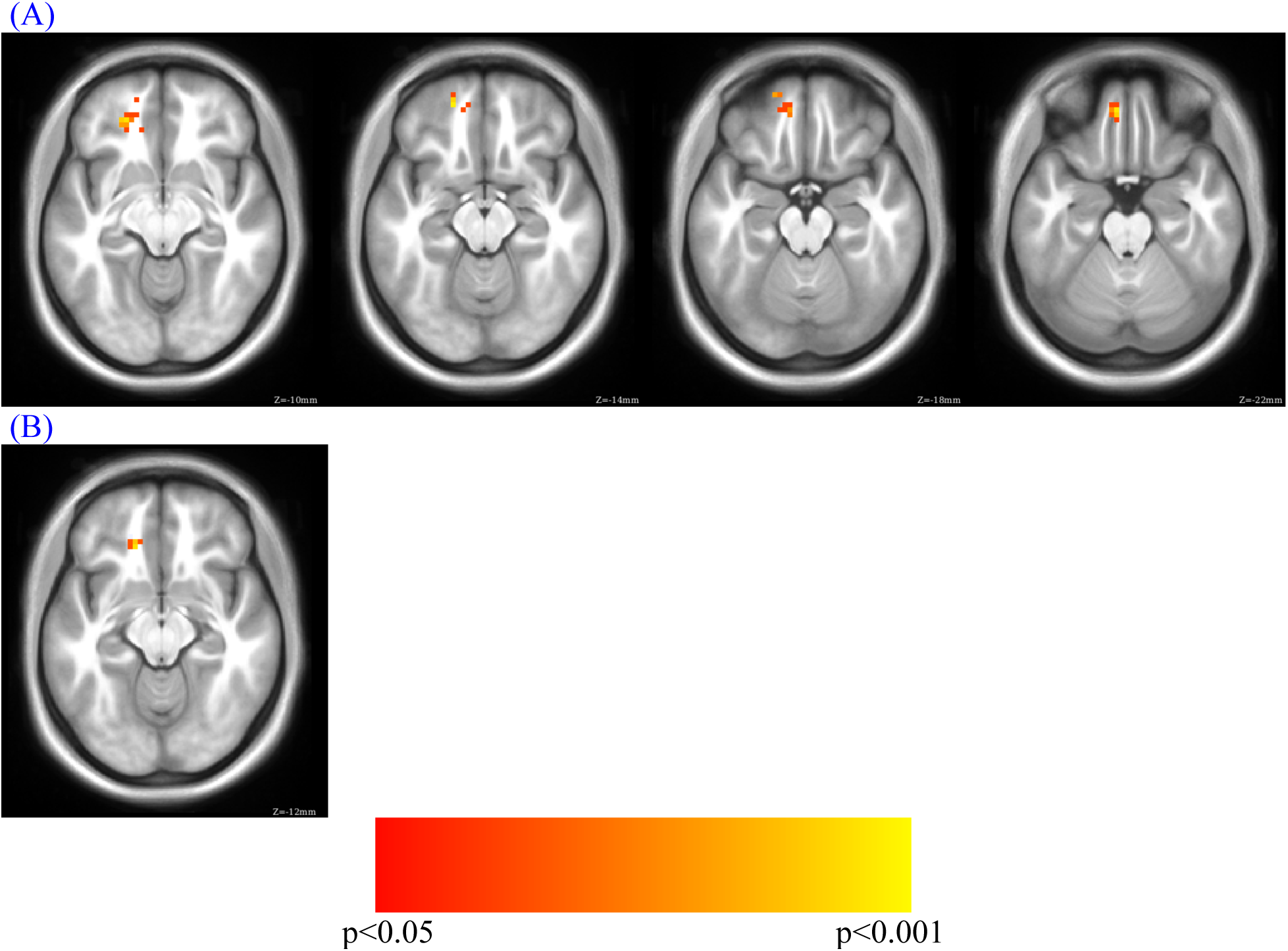
PHQ-8 regression for sgCC connectivity seedmaps for A) Cohort 1 (MNI Z of -10 to -22 in steps of 4) and B) overlap between Cohort 1 and Cohort 2 (MNI Z of -12). Cohort 1 PHQ-8 results were masked with the volume-corrected cluster identified in the Cohort 1 phone usage analysis (unlock duration) and Cohort 2 PHQ-8 results were masked with the PHQ-8 results from Cohort 1.

**Table 3.** Results for the correlation of sgCC RSFC seedmaps, correlated with PHQ-8, masked by phone screen time results (Top). Overlap between Cohort 1 and Cohort 2 for sgCC RSFC seedmaps correlated with PHQ-8 (Bottom). Subpeaks are at least 8mm apart. * signifies that cluster didn’t pass volume correction. Cohort 1 PHQ-8 results were masked with the cluster identified in the Cohort 1 phone usage analysis (unlock duration) and Cohort 2 PHQ-8 results were masked with the PHQ-8 results from Cohort 1.

| **Cohort 1** |  |  |  |  |  |
| --- | --- | --- | --- | --- | --- |
| **Best Estimate of Region** | **X** | **Y** | **Z** | **T** | **Extent** |
| Left OFC | -21 | 42 | -12 | 3.19 | 63 |
|  | -18 | 51 | -15 | 3.09 | Subpeak |
|  | -6 | 48 | -21 | 3.04 | Subpeak |
| Right OFC* | 24 | 51 | -9 | 2.55 | 15 |
|  | 18 | 42 | -12 | 2.19 | Subpeak |
| **Overlap Between Cohorts** |  |  |  |  |  |
| **Best Estimate of Region** | **X** | **Y** | **Z** | **T** | **Extent** |
| Left OFC | -15 | 33 | -12 | 2.98 | 8 |

PHQ-4 EMAs correlated with sgCC connectivity maps identified peaks in Cohort 1 and 2, but there was no overlap in the clusters between the Cohorts (Table S1). In Cohort 1 no significant clusters were identified when PHQ-4 was masked with Cohort 1 unlock duration. As Cohort 1 didn’t identify any regions which passed volume-correction, there was no overlap of significant volume-corrected regions between Cohort 1 and Cohort 2 for PHQ-2. As such, Cohort 2 results were masked with the Cohort 1 unlock duration cluster which identified one significant cluster with the peak at -15, 30, -12 (peak T = 3.71, voxel extent = 41, volume corrected to p < 0.05). Two clusters were identified that didn’t pass volume correction were also identified at -9, 51, -18 (peak T = 2.87, voxel extent = 28, volume correction *ns*) and 24, 39, -15 (peak T = 1.87, voxel extent = 9, volume correction *ns*).

As PHQ-4 includes two anxiety questions, we wanted to examine if there were more robust results when restricting the analysis to the two depressive symptoms, specifically PHQ-2. As Cohort 1 didn’t identify any regions which passed volume-correction, there was no overlap of significant volume-corrected regions between Cohort 1 and Cohort 2 for PHQ-2. As such, Cohort 2 results were masked with the Cohort 1 Unlock Duration cluster which identified 1 cluster which passed volume correction, with the peak at -18, 30, -12 (peak T = 3.81, voxel extent = 60, volume corrected to p < 0.05). One cluster was identified that didn’t pass volume correction with peak at -9, 42, -27 (peak T = 2.93, voxel extent = 40, *ns*).

### Overlap across analyses here

Given the similarity of regions found across the PHQ analyses in Cohort 2, we investigated the overlap between the results of PHQ 2/4 masked by the Cohort 1 unlock duration, with 39 voxels out of the 41 voxels identified in the PHQ-2 analysis overlapping with the PHQ-4 analysis. The overlap between Cohort 2 PHQ-2, 4 and 8 identified 11 voxels, which are located around the peaks of the PHQ-8 analysis.

## Discussion

In the current manuscript, we provide an example of how to link passive smartphone metrics, active smartphone-based surveys of mental health and computer-based surveys of mental health with brain connectivity measures. Specifically, RSFC between the subgenual cingulate cortex, a region previously implicated in depression, and nearby ventral prefrontal regions, was strongly related to unlock duration, such that more connectivity was associated with more screen time, which has been implicated as being related to self-reported depressive symptoms. The link between RSFC and individual differences has long been established but extending that and combining it with an individual’s behavior inferred from smartphone sensors provides exciting new directions. While the results presented here are a relatively simple analysis of complex, highly dimensional data, we discuss some methods which could be used in the future to combine these highly multivariate and complex datasets in exciting ways.

Phone-related screen time, which we define here as the amount of time a phone is unlocked, or unlock duration, has previously been shown to be related to self-reported depression levels (Twenge et al., 2018; Wang et al., 2018). An exploratory analysis in Cohort 1 of the correlation between unlock duration and sgCC seedmaps identified a large cluster which extended from the anterior caudate to medial frontal orbitofrontal cortex and dorsally to medial prefrontal cortex, a result which was replicated in Cohort 2 with a smaller voxel extent, even though the sampling rate for screen time was greatly reduced, reducing our sensitivity to pick up individual differences in phone usage for this cohort. Next, to determine if depressive symptoms showed a similar pattern of connectivity between sgCC and ventral prefrontal cortex the cluster from Cohort 1’s unlock duration analysis was used as a mask with PHQ-8, a commonly used survey to assess depressive symptoms in the general population. Two small clusters of overlap were identified in the left OFC, one of them neighboring voxels that were identified to replicate in the unlock duration analysis between the Cohorts. While these clusters are not large and would not necessarily survive volume correction on their own, observing similar regions across Cohorts and analyses suggests that there is a link between depressive symptoms and related behaviors and sgCC-OFC connectivity, particularly left OFC that should be further investigated. The PHQ-4, which contains two depression questions and two anxiety questions, did not show the same robust relationship across both Cohorts, with no voxels overlapping, although Cohort 2 identified a cluster in the left OFC which overlapped with results observed with PHQ-8 in both Cohorts. Connectivity between the sgCC seed (BA 25), located at 0, 25, -10 and the left OFC region around -15, 33, -12 shows a consistent relationship between self-reported depressive symptoms and screen time, which has previously been associated with depression. Increased connectivity between sgCC, a region involved in processing of valenced information about the self (Moran et al., 2004) and OFC, which is involved in valuation and reward processing has been linked increased depressive symptoms and *screen time* across both Cohorts. Similar results were observed with PHQ-2, which only contains the two questions directly related to mood. It seems quite plausible that regions involved in valence processing related to the concept of self and a more general reward valuation processing region would have increased connectivity in individuals with higher depressive symptoms.

We have shown that resting-state functional connectivity of the brain, as measured with MRI, in two separate Cohorts of individuals, with two separate MRI’s and two separate versions of the StudentLife applications show similarity in the results observed. The cluster identified with the unlock duration analysis covered an extent similar to that of the limbic network previously identified (Choi, Yeo, & Buckner, 2012; Yeo et al., 2011). Due to the constraints we imposed on the analysis, all of the subsequent results were within this area, but noticeably, many of the results were proximal to the left OFC, which is also a member of a set of nodes which are commonly activated during reward processing and can form their own preferentially coupled system (Huckins et al., 2018) and is identified as a peak of the term “reward” in reverse-inference meta-analyses using Neurosynth (Yarkoni et al., 2011).

## Limitations and Future Directions

The current work is a first-pass at analyzing longitudinal multi-cohort, multimodality data and has several limitations. There are several ways in which future research may provide a more comprehensive survey of the relationships between the diverse set of features provided from passive smartphone sensing, functional brain connectivity measures and self-reported measures of depression or other mental health metrics. The relatively small number of clinically depressed individual in the current sample weighs the results heavily on the RSFC and passive-sensing features from those individuals. Test-retest within the moderately sized samples allows for identification of factors with reliable cross-cohort replicability in RSFC both and passive-sensing features. Ideally, similar sensing features could be collected across many sites, allow for identification and characterization of depressive subtypes that span across passive-sensing and RSFC as has been done by Drysdale and colleagues (2016) with RSFC and survey data. In the current study, particularly Cohort 2 in which data quality was actively monitored, we retained a relatively large portion of individuals from those scanned (see Table 1). The sample sizes used here would have been considered relatively large several years ago. Increased sample sizes in the current study would help future analyses given the large number of features from both passive mobile smartphone sensing and RSFC. An outstanding question is if long-term changes in depressive symptoms can be better predicted by RSFC or smartphone sensing metrics at the initiation of the study or if changes in either of these over time parallel depressive symptoms. Ideally to assess this a large number of individuals would be tracked over multiple years. In the second Cohort our working group aims to track them over multiple years while eventually increasing the number of individuals enrolled. Furthermore, including multiple sites, as the ABCD study does (Volkow et al., 2017), would increase applicability to a wider population. Multiple research sites are currently collecting MRI data, self-reported surveys and smartphone sensing metrics. An unresolved issue is what, exactly, is the optimal approach to analyze the huge amounts of multivariate data produced by these methods.

### Application changes Between cohorts

In the current study, unlock duration data collection changed between the cohorts. In Cohort 1, unlock duration was continually sampled, while in Cohort 2 unlock duration was adaptively sampled between 10 and 30% of the time. This change was instituted to optimize battery life, a primary limitation to users being willing to keep the StudentLife app on their phone. By decreasing the amount of time sampled from 100% to 10-30%, our ability to accurately estimate unlock duration may decrease slightly as evidenced by an observed decrease in peak effect (T-value) and voxel extent. As with all passive and active smartphone features, the ability to collect data must be weighed against the invasiveness to the user experience, either through app prompts or decreased battery life and phone speed.

### Temporal factors related to school

The demands of the academic term provide a generally applicable path of stress which is shaped over the term. Avoiding, or potentially purposefully collecting MRI data during finals, which may be particularly stressful, or during popular social weekends may lead to changes in stress levels, sleep patterns and other variables which could alter connectivity patterns and self-reported behavioral data that would have otherwise been observed. In the study herein, we attempted to scan before finals and avoid well-known “party weekends”. Future studies may be able to capitalize on temporal differences in stress and depression levels by scanning at these peak times of stress or sleep deprivation and comparing that data to less stressful times, such as the beginning of the term.

### Functional Differences and Alignment Across Individuals

Resting-state functional connectivity shows robust and relatively reliable connectivity across large groups of individuals across methods (Gordon et al., 2016; Yeo et al., 2011). Meanwhile there are individual differences in the cortical extent of large-scale functional regions across individuals and even the network membership of these regions can vary (Gordon et al., 2017). Furthermore, critical to identifying group and individual differences is acquiring a large quantity of high-quality data (Gratton et al., 2018). Defining networks on an individual basis will likely help in the pursuit of the individual differences in brain connectivity that underlie depression. Variability in resting-state functional connectivity has been observed at the functional parcel level, but what about at finer resolutions? While a departure of traditional anatomical alignment methods, hyperalignment is a method which attempts to align brain based on similar response patterns in high-dimensional space (Guntupalli et al., 2016). While this method originated using time-locked dynamic stimuli such as a movie, it has recently been applied to resting-state functional connectivity as connectivity hyperalignment (CHA), which revealed both coarse-scale, areal structure as previously observed, along with fine-scale structure which was previously inaccessible. Applying connectivity hyperalignment to RSFC data is will hopefully allow for increased ability to discern individual differences in depression and other mental-health metrics.

### Voxelwise Resting-State Functional Connectivity

A relatively simple first-pass method is to target specific region and feature pairs. If there are *a priori* hypotheses related to the topic of interest it may be possible to look at connectivity from one region using seed maps or between a small number of regions and relate them to specific passive-sensing features. As shown here this is plausible but even correlating seed maps with 1 sensing variable leads to potential multiple comparisons issues based on the 50,000+ voxels in the brain using a 3mm^3^ voxel size. Recent statistical simulations have suggested an increased false-positive rate associated with older versions of 3dClustSim, a function of AFNI (Cox, Chen, Glen, Reynolds, & Taylor, 2017). Indeed, the authors of 3dClustSim now suggest using a different algorithm with the same program, the autocorrelation function (ACF) with a high p-value threshold per voxel to minimize the possibility of false-positives. In some datasets, at lower p-value thresholds ACF requires a much larger voxel-extent than the old version of 3dClustSim. The increased voxel-extent may make it less likely to identify smaller functional regions in a whole-brain regression using a lower per-voxel p-value threshold (p<0.05). This evolution of methods decreases the rate of the false-positives which is critical but requires a larger expected functional region, a very strong effect size or a very large number of participants. Across all possible methods presented here there are a variety of factors which should be taken into consideration to decrease false positive rates. Having a large number of subjects to draw data will increase the portion of the population sampled.

If possible having two distinct Cohorts to analyze then looking for overlap in results between the Cohorts would decrease false positives due to random sampling, Cohort specific variance, and further increase the total size of individuals sampled. The above factors apply to most any study. With passive smartphone mobile sensing there are many features which can be measured or computed based on the intersection of multiple features. For example, “phone unlock duration” is a very simple metric, which measures the time that the smartphone was unlocked. This can be further broken down into location specific features, such as “phone unlock duration at dorm” or “phone unlock duration at study places” by looking at the intersection of location on a geo-tagged campus and “phone unlock duration. Given the large number of initial features that can be calculated, along with the nearly endless number of meta-features that could potentially be generated, making sure that the feature is relatively straightforward to calculate and interpret should be at the forefront of anyone analyzing passive-mobile phone sensing features. Features that are difficult to calculate or interpret could easily be embedded with unforeseen confounds. Furthermore, such features should be validated to make sure they are measuring the effect or phenomena they are supposed to in an accurate manner.

Typically, only features with sensing data from many days should be used to get a more stable estimate of that features’ value. While putting a sensing application of many students’ phones may seem like a plausible method for maximizing data collect, there are a variety of factors which can lead to reduced data collection, potentially rendering an individual’s sensing data unusable. Phone operating system (OS) updates can often change application permission or render the sensing application completely useless. To avoid this beta testing should be done as early as possible and new versions of the application that are compatible with the latest OS pushed to participants. Participant non-compliance or attrition is another important factor to consider. Individuals may delete the application, limit its permissions within the OS or otherwise limit the researcher’s’ ability to accurately measure data. Clearly, it is the individual’s choice to continue to participate in any study, particularly one where large amounts of data are being collected (anonymously) on their habits. It may be difficult for the researcher to determine if the individual has deleted the application or simply not uploaded their data in while. Finally, a rate of attrition is expected in all longitudinal studies and some individuals may simply decide that they do not wish to continue their participation in the study.

### Whole-brain and network-based connectivity

A possible method to deal with the large number of comparisons related to voxelwise or whole-brain connectivity is to simply look at connectivity between a set of predefined regions or parcellation (Huckins et al., 2018; Gordon et al., 2016; Poldrack et al., 2015; Power et al., 2011; Yeo et al., 2011). Connectivity between each pair of regions can be correlated with the sensing feature of interest. Unfortunately, many of the commonly used parcellations have many nodes, which increases the total number of comparisons in a nonlinear manner as the number of nodes increases. The number of comparisons can soon approach the number of comparisons evident when using voxelwise seed maps without methods such as voxel extent to appropriately correct for the associated multiple comparisons.

A simple but perhaps relatively unsophisticated sophisticated method is to calculate mean connectivity within a functional system or network. The system or network would be determined off of data driven approach such community detection using a random walk technique like InfoMap (Rosvall & Bergstrom, 2008) or regions identified as being part of a coherent functional system using another method. In this approach, the mean of all Fisher r-to-z transformed correlation values between nodes of interest is calculated. For example, mean connectivity within the Cingulo-Opercular network would be calculated between all nodes or parcels belonging to that network. Between-network or system connectivity can also be calculated by taking the mean of all pairwise connections between the two networks of interest. This can greatly reduce the number of total connections observed, thus reducing the multiple comparisons problem mentioned under the whole-brain connectivity section. One drawback to this method is that it is not selective about which connections it is using in the calculation - specifically, that it may be and probably is including connections that are not physiologically or psychologically relevant.

A plausible may to reduce the number of connections by selections ones that are likely to be “real”, such that information may actually travel through that connection on the neural level, even if not on a first-order or even second-order synapse. Multiple approaches have been taken to identify meaningful connections. Within or between networks there are likely to be positive and negative correlations, which then somewhat cancel out. One could take the absolute value of each connection before averaging across the network, but this would introduce bias in any connections with a distribution of correlation values that included positive and negative values. Values of correlation, or connectivity measures in the brain vary by orders of magnitude. Identifying a multiscale network backbone that accounts for important connections within and between communities, regardless of the connectivity strength would be a method to decrease the number of connections analyzed. One way of identifying the network backbone is to use the z-value from each connection as the weight, or amount of information that could travel between the two brain regions that the connectivity was estimated from. A group did just this (Serrano, Boguñá, & Vespignani, 2009), identifying connections which are statistically relevant across multiple scales of connectivity, work which has been extended non-parametrically (Foti, Hughes, & Rockmore, 2011). By identifying the network backbone for each individual, it may be plausible to identify a variety of subcategories or continuums of depression along which different symptom severities fall for each individual, along with passive smartphone monitoring will allow for greater insight into interactions of behavioral, self-report and physiological RDoC matrix criteria.

### Wrangling high dimensional data

A variety of techniques can be used to extract information from data that are both longitudinal and high-dimensional; that is, situations where the data are collected from participants at multiple time points and the number of covariates begins to approach, or even surpasses the number of subjects in the dataset (Cheng, Honda, Li, & Peng, 2014; Chu, Li, & Reimherr, 2016; L. Wang, Zhou, & Qu, 2012; Zipunnikov et al., 2014).

As has been mentioned repeatedly above, both with resting-state and passive smartphone sensing there are a large quantity of features and analyses that can be generated. In the current study we chose features that were reasonable based on previous data but are unlikely to be the optimal features that describe the relationship between depression, passive mobile sensing and brain connectivity. Multiple approaches could be taken with data from both sources. One approach which would greatly decrease the number of features that were necessary including trying to create a singular propensity metric, or biomarker of depression for both the resting-state fMRI data and a separate one for the sensing data then observing the relationship between the two. Alternatively, data reduction techniques such as independent component analysis could be applied to each group then the relationship between them could be measured. Many researchers have taken a “risk” or “propensity” score approach, where they generate models which contain predictive variables (gender, substance use, family history) pertinent to the outcome of interest and use the propensity score as a regressor when doing analyses at the group or individual difference level (Hansen et al., 2012; Stuart, 2010). This could be applied to smartphone data, but only once appropriate sensor features, and model have been calculated. By creating a unitary risk feature multiple comparison issues can be greatly mitigated. Data reduction techniques that account for variance that is common between two data modalities such as joint ICA, parallel ICA and CCA-Joint ICA, which has been implemented for combining high-dimension data across fMRI and genetic data (FusionICA, available from http://mialab.mrn.org/software/fit/).

### Unresolved Questions about Directionality and Timing

In the current sample, resting-state fMRI data is from 1 time-point while mobile smartphone sensing data is dynamic and collects data over a longer period of time. An unresolved question is if changes in fMRI data across multiple sessions reflects or predicts changes in smartphone usage. Likely a more sensitive measure would be to do the reverse - using changes in smartphone usage, which is continuously monitored, to predict when there may be changes in brain connectivity as measured by fMRI. Changes in depressive symptoms have been successfully predicted with passive smartphone features (Wang et al., 2018), and may be useful for signaling when an individual should be referred to clinical services or brought in for a subsequent fMRI session. Longitudinal penalized functional regression is a method designed to deal with multiple timepoints of both exposure and outcomes (Goldsmith et al., 2012) which may help provide insight into the temporal association between brain connectivity, depression and phone usage.

### Moderating Factors of RSFC

RSFC has repeatedly been shown to be relatively stable across individuals and time, displaying similar network structure across thousands of individuals. While similar network structure and connectivity patterns are observed between sites, preprocessing methods, and Cohorts, differences between individuals are observed across individual differences in personality, affect and current mood have been related to alterations in RSFC. Furthermore, individual differences in the network structure on an individual level have been observed. Properly mapping individual differences in networks across the cortex would allow for better cross-subject alignment. The network assignment of particular regions may in itself be linked to depressive symptoms, while lining up networks would allow for the proper comparison of networks across individuals. Additionally, the current state physiological state an individual is in, such as food satiety or caffeination status can influence their mood (Rogers & Lloyd, 1994) and has also been shown to influence an individual’s brain connectivity (Poldrack et al., 2015). While there are a variety of factors that can influence RSFC, reliable individual differences across brain disorders have been observed in previous studies and here. As the predictive accuracy of RSFC or other neuroimaging methods increases the field may move closer to using MRI as a biomarker of depression, as has been done with physical pain (Atlas, Bolger, Lindquist, & Wager, 2010; Wager et al., 2013).

## Conclusions

In summary, we have identified proof-of-concept relationships between resting-state functional connectivity of the brain, web-based self-reported surveys of depressive symptoms (PHQ-8), a passive mobile smartphone sensing feature (unlock duration) and mobile smartphone based ecological momentary assessments of depressive symptoms (PHQ-4). An important mental health implication is that the amount of time spent using a phone is correlated with depressive symptoms. Further, these symptoms, both before time-of-scanning (PHQ-8) and after time-of-scanning (PHQ-2/4), show a relationship with connectivity between areas implicated in depression, reward and processing of valenced self-relevant material. Importantly, these initial results predominantly replicate across the two separate cohorts, increasing the applicability and scope of the findings herein. Although the current results do not elucidate causality in the relationship between screen-time, depression and brain connectivity, future work should aim to do so, especially given recent changes to public policy, with professional groups such as the American Academy of Pediatrics providing suggesting screen-time limits and policy and investor groups calling on media device makes such as Apple and other phone makers. In the current work we extend previous research, replicate results across multiple MRI scanners and cohorts all while combining data from a while variety of sources. The analyses done here are by no means comprehensive and we hope that the findings of this study and future research methods proposed herein are useful to a wide-range of researchers. Ultimately continuation and extensions of this research has the potential to provide important insights into mental health, as well as inform psychological treatments and other interventions.

## Funding

NIMH #’s 5R01MH059282-12 and 5R01DA022582-10.

## Acknowledgements

We would like to acknowledge Yaroslav O. Halchenko for assistance with MRI data conversion and BIDS organization and Terry Sackett for assistance with MRI data collection.

## Supplemental Material

**Figure S1.**
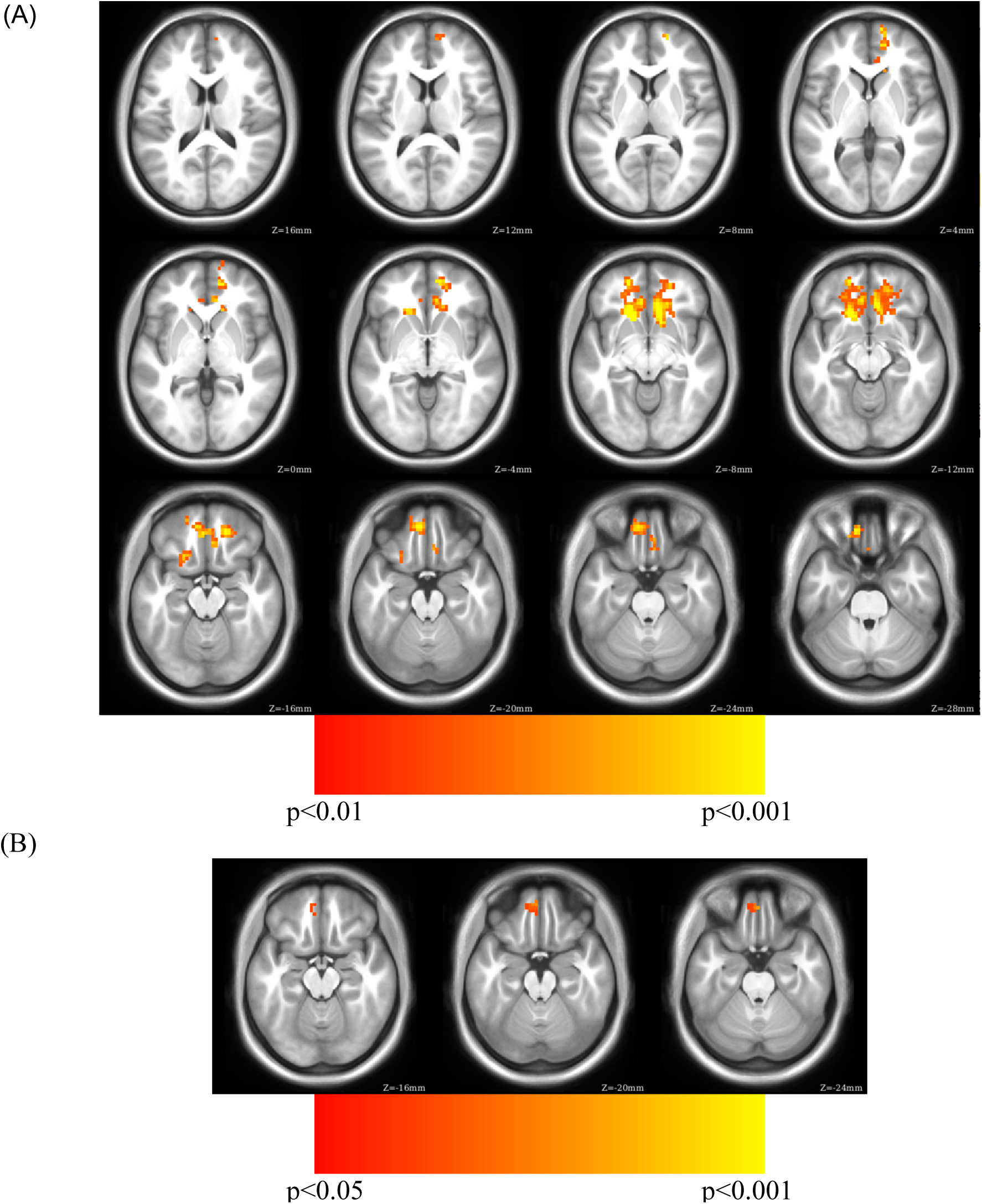
Exploratory analysis correlating sgCC RSFC seedmaps correlated with mean unlock duration in Cohort 1 (A) identified a cluster with a positive relationship to unlock duration in the ventromedial prefrontal cortex (p<0.01, volume corrected using ACF to 0.001, k>449) shown on axial slices. Replication of the relationship between unlock duration and sgCC connectivity between Cohorts 1 and 2 are observed in (B). The cluster identified in Cohort 1 showed a positive relationship with unlock duration was subsequently used to restrict the area interrogated in Cohort 2, showing positive clusters in ventral medial orbitofrontal cortex (p<0.05, volume corrected using ACF to p<0.05, k > 38).

**Figure S2.**
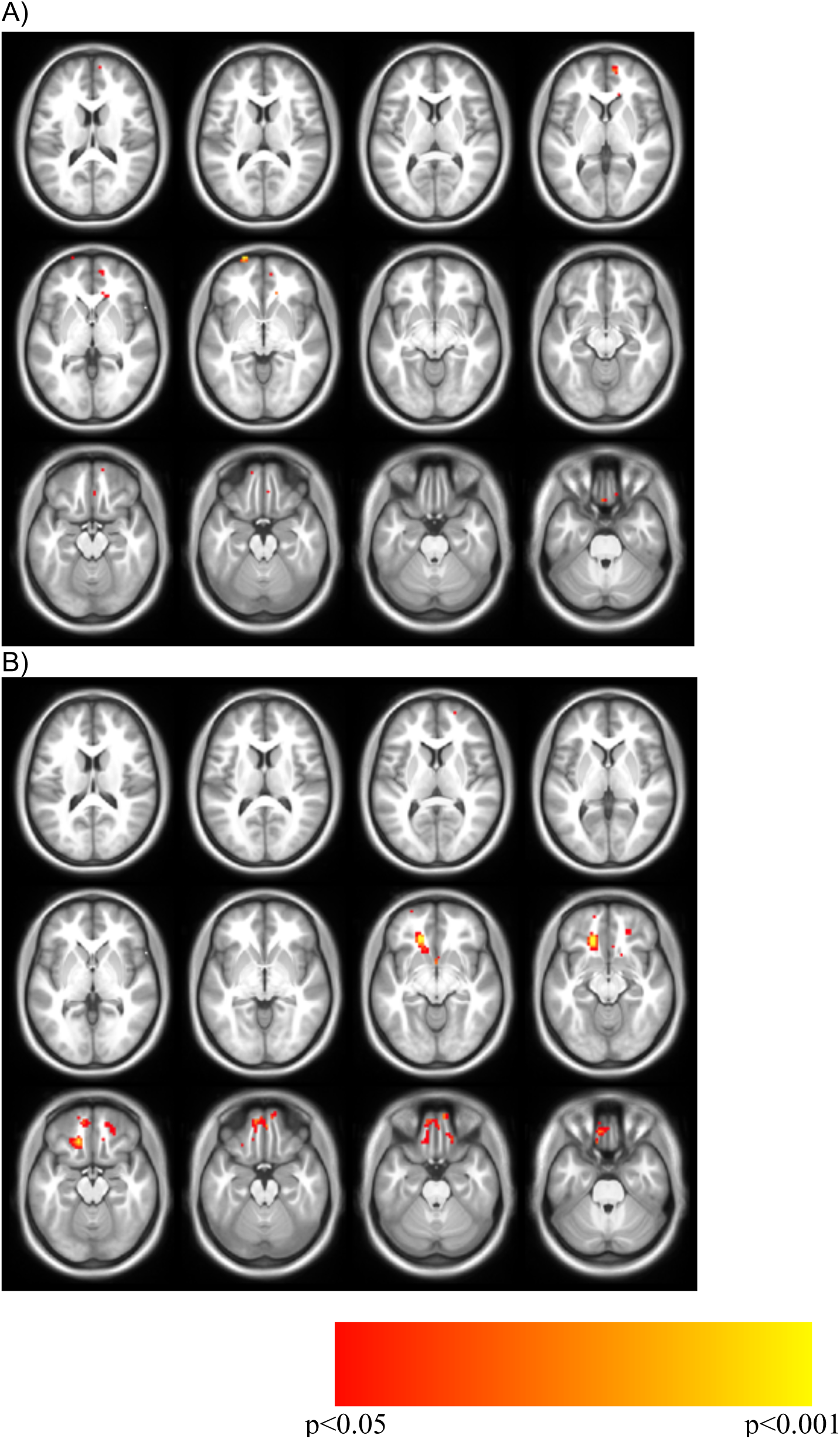
PHQ-4 regression for sgCC connectivity seedmaps for A) Cohort 1, B) Cohort 2 masked by the results from unlock duration in Cohort 1. No common regions were found between the two analyses.

**Figure S3.**
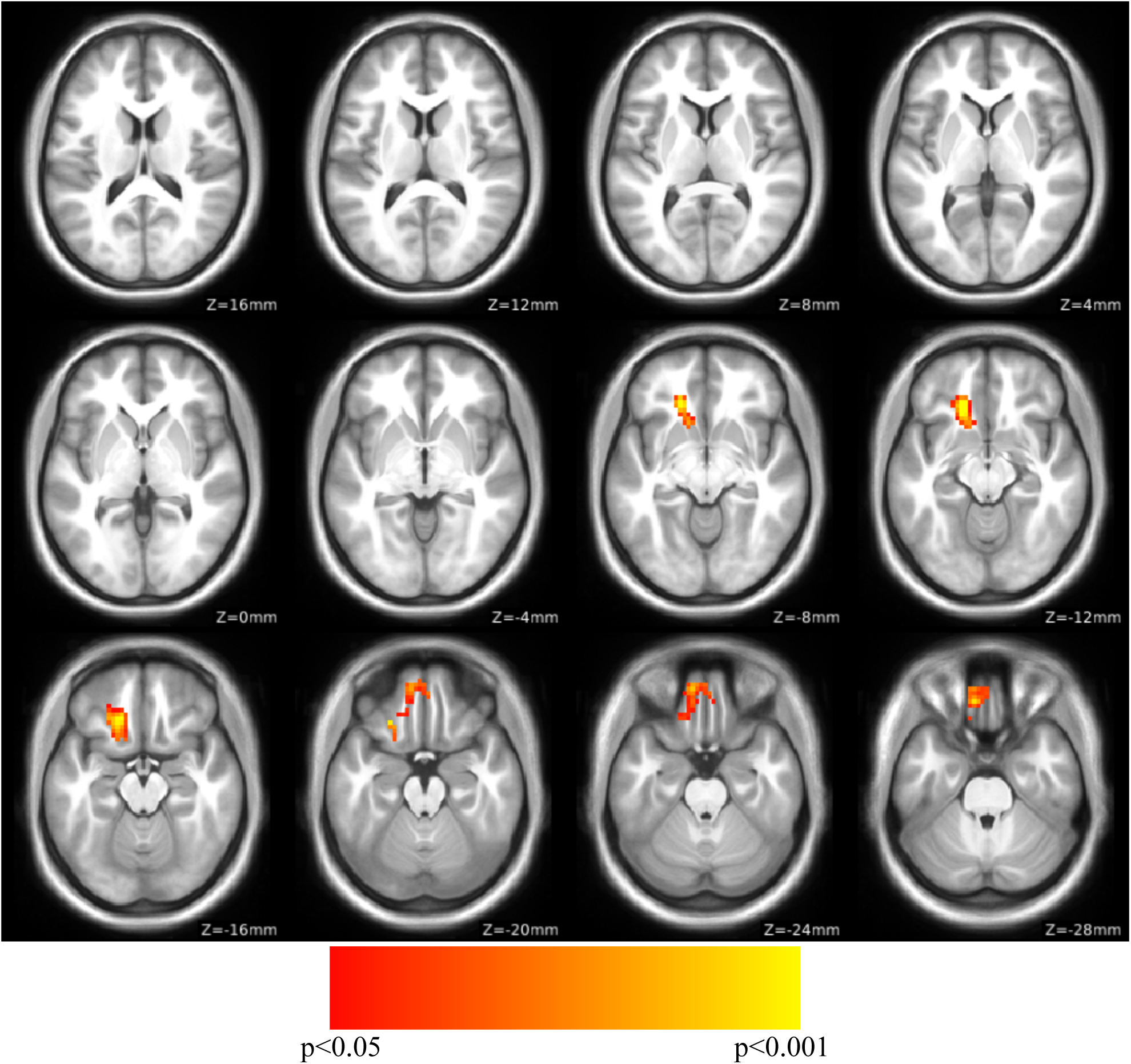
PHQ-2 regression for sgCC connectivity seedmaps for Cohort 2 masked by the results from unlock duration in Cohort 1, volume corrected to p < 0.05. No significant cluster were found in the Cohort 1 analysis and thus there was no overlap between the two analyses.

**Table S1.** Exploratory analysis identifying peaks with a positive relationship from sgCC RSFC seedmaps correlated with phone-based surveys of depressive symptoms (EMA form of PHQ-4). Only clusters showing overlap with the Cohort 1 unlock duration analysis, with a positive relationship between PHQ-4 and connectivity with greater than 5 contiguous voxels are reported here. * signifies that cluster didn’t pass volume correction.

|  |  |  |  |  |  |
| --- | --- | --- | --- | --- | --- |
| **Cohort 1** |  |  |  |  |  |
| **Best Estimate of Region** | **X** | **Y** | **Z** | **T** | **Extent** |
| Superior Frontal Gyrus* | -21 | 72 | -3 | 2.99 | 7 |
| Medial PFC* | 9 | 57 | 3 | 2.31 | 7 |
| **Cohort 2** |  |  |  |  |  |
| **Best Estimate of Region** | **X** | **Y** | **Z** | **T** | **Extent** |
| Ventromedial PFC | -15 | 30 | -12 | 3.71 | 139 |
|  | -9 | 51 | -18 | 2.87 | Subpeak |
|  | 3 | 48 | -21 | 2.49 | Subpeak |
| Anterior Medial OFC* | 9 | 57 | -24 | 2.38 | 7 |
| Right OFC* | 18 | 33 | -24 | 2.11 | 5 |
| Right OFC* | 24 | 39 | -15 | 1.87 | 13 |
|  | 15 | 51 | -15 | 1.72 | Subpeak |
| **No Cohort Overlap Observed** |  |  |  |  |  |

